# Effects of Alternative Splicing-Specific Knockdown of Tjp1 α+ by Rbm47 on Tight Junctions Assembly during Blastocyst Development

**DOI:** 10.1101/2023.07.18.549609

**Authors:** Jiyeon Jeong, Inchul Choi

**Affiliations:** Division of Animal and Dairy Sciences, College of Agriculture and Life Sciences, Chungnam National University, Republic of Korea

**Keywords:** Rbm47, ZO-1 alpha, TJP1, Tight junctions, Blastocyst

## Abstract

Tjp1 α+ is considered a crucial protein involved in the stepwise assembly of tight junctions (TJs) between compaction and blastocoel cavitation in early development. In this study, we investigated the specific role of Tjp1 α+ in TJ formation by employing an alternative splicing-specific knockdown of the Tjp1 α+ exon. To deplete Tjp1 α+ expression, we used siRNA targeting RNA-binding protein 47 (Rbm47), which induces the inclusion of the α+ exon in Tjp1 mRNA. The knockdown resulted in approximately 85% reduction in Rbm47 mRNA levels and 75% reduction in Tjp1 α+ mRNA levels in blastocysts. Surprisingly, despite this knockdown, blastocyst development and TJ permeability of trophectoderm were unaffected. Additionally, we observed an interaction between Tjp1 α- and Ocln in Rbm47 knockdown blastocysts, suggesting a compensatory role of Tjp1 α-. Overall, our findings indicate that Tjp1 α+ is not essential for the stepwise assembly of TJs and the completion of TJ biogenesis during blastocyst development in mice although a minimal amount of remaining Tjp1 α+ is sufficient for TJs assembly.

**Summary statement:** Selective loss of Tjp1 α+ mediated by Rbm47 knockdown did affect mouse blastocyst development, suggesting that Tjp1 α+ may not be crucial for stepwise TJs assembly during blastocyst development

## Introduction

During early embryonic development, the fertilized embryo undergoes rapid cell division and develops into a blastocyst, which consists of two distinct cell lineages, the inner cell mass (ICM) and the outer trophectoderm (TE) layer, surrounding a fluid-filled cavity known as the blastocoel. The formation, maintenance, and expansion of the blastocoel rely on the accumulation of water, which is mediated by ion gradients (Na/K-ATPase) and water channels (aquaporins) across the TE layer, as well as the assembly of tight junctions (TJs) on the apical and basolateral membranes of the outer cell layer (Cockburn and Rossant, 2010; Eckert and Fleming, 2008; Watson and Barcroft, 2001).

Several tight junction-associated transmembrane proteins, such as OCLN, CLDN4, CLDN6, and JAM1, as well as cytoplasmic adaptor proteins PARD6B, TJP1, and TJP2, are essential for blastocoel formation and expansion during the early stages of embryonic development (Alarcon, 2010; Katsuno et al., 2008; Moriwaki et al., 2007; Saitou et al., 2000; Sheth et al., 2008). The transcription factor AP-2 gamma (Tfap2c) regulates the expression of these genes during the morula-to-blastocyst transition (Choi et al., 2012; Lee et al., 2016). In addition, TJ-associated proteins, including CXADR, ADAM10, and CPEB2, are necessary for TJ assembly during blastocyst formation in pig and mouse embryos (Jeong et al., 2022; Jeong et al., 2019; Kwon et al., 2016a; Kwon et al., 2016b).

The assembly of TJs is a dynamic process that occurs in a stepwise sequence between compaction and blastocoel cavitation. The assembly of peripheral membrane scaffold protein ZO-1(also known as TJP1), together with rab-GTPase, rab13, takes place immediately upon compaction, followed by the assembly of peripheral membrane proteins cingulin and ZO-2 (also known as TJP2) during the 16-cell stage. Finally, during the 32-cell stage, late expression of ZO-1α+ and its assembly with the transmembrane proteins OCLN and CLDN-1/3 generate the paracellular seal between TE cells during blastocyst formation. ZO-1α+ is critical in this biogenesis process (Eckert and Fleming, 2008; Sheth et al., 1997; Sheth et al., 2000a).

In our recent publication (Jeong et al., 2022), we presented findings on a post-transcriptional mechanism regulated by CPEB2, an mRNA-binding protein, which plays a vital role in determining the subcellular localization and stability of Tjp1 mRNA during tight junctions biogenesis in blastocyst development. Building upon these significant findings, our current study aims to investigate the biological function of a specific isoform of TJP1, known as ZO-1α+, in the context of tight junction assembly during mouse blastocyst formation. To accomplish this, we utilized siRNA targeting RNA-binding motif protein 47 (RBM47) to selectively disrupt the alternative splicing of TJP1, specifically eliminating the ZO-1α+ isoform. (Kim et al., 2019). This approach allows us to assess the impact of the absence of the ZO-1α+ isoform on tight junction assembly dynamics.

## Results and Discussion

In this study, we examined the function of Tjp1 variant, Tjp 1 α+ by alternative splicing specific gene knock-down by siRNA of Rbm 47 during TJs assembly in mouse blastocyst development. We first designed Tjp1 variants specific primer sets to distinguish between the two variants of Tjp1 and a common primer set that amplifies the common region of Tjp1. The design of alternative splicing specific primer sets that include exon 20 (α+) and extended exon 27 (α-) are shown in Figure 1. The expression patterns of each variant of Tjp1 were consistent with those reported in a previous study (Sheth et al., 1997). Tjp1 α-was detected throughout preimplantation development, from 1-cell to blastocyst. Upregulation was particularly observed from the 8-cell stage onwards. Tjp1 α+ was highly expressed from the morula stage onwards (Figure 1B and C).

**Figure 1.**
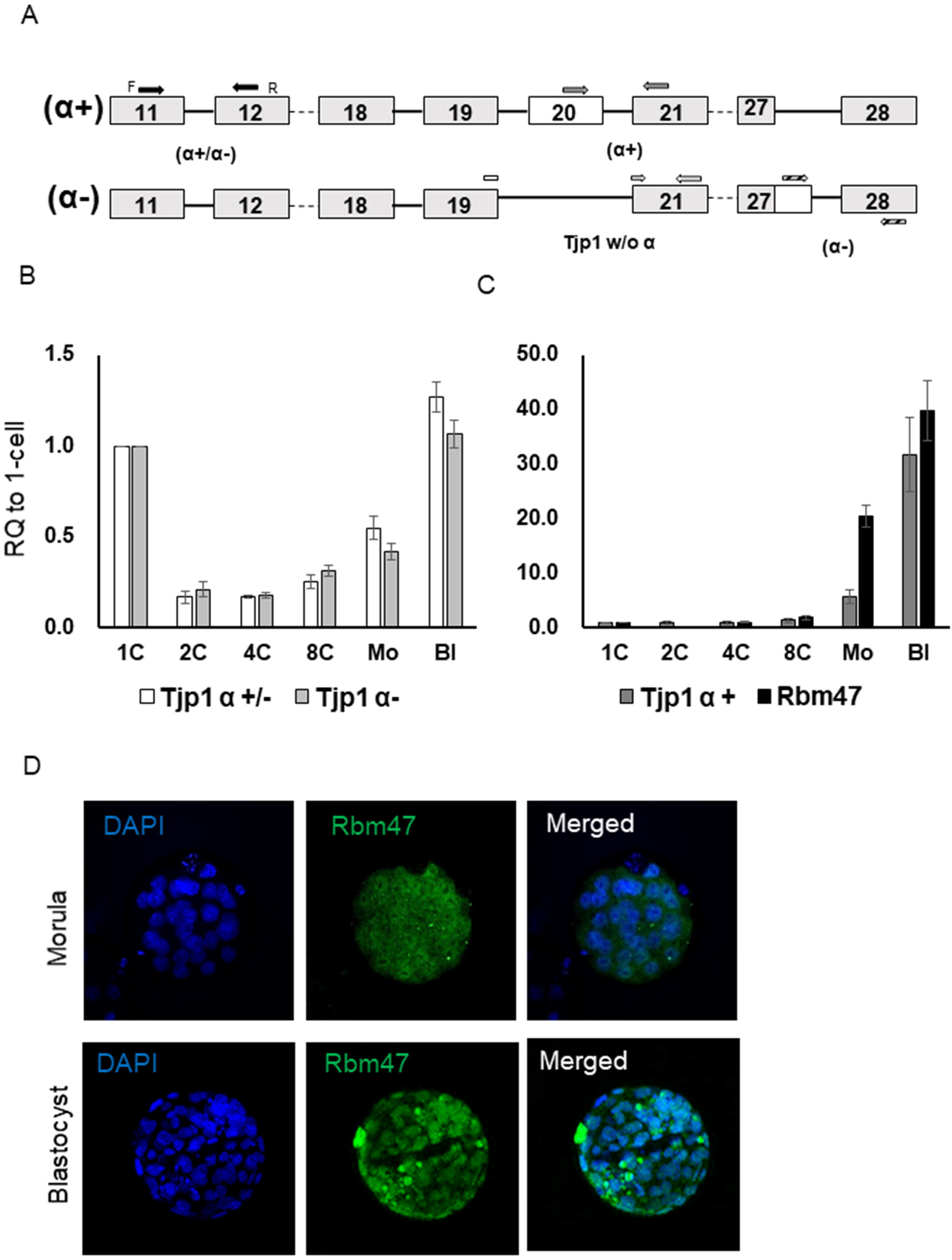
Expression analysis of TJP1 isoforms and Rb47. (A) The position of both isoforms of Tjp1 (exon11/12; α+/α-), the α-domain (exon20/21) and two Tjp1 α-specific (exon 19/21; w/o α and alternatively spliced exon 27/28; α-) primers used for cDNA amplification of Tjp1 isoforms. Arrows mark the position and direction of primers used for cDNA synthesis. Transcription levels of Tjp1 α+/-, Tjp1 α variant (B), and Tjp1 α+/-, Tjp1 α-, Rbm47 (C). Transcription levels of Tjp1 α+/α-, Tjp1 α-, and Rbm47, shown as relative quantification (RQ). Error bars represent mean ± standard error.

In this study, we examined the expression of RNA-binding protein 47 (Rbm47), a splicing regulator known that induces the inclusion of the α+ exon in Tjp1 mRNA (Kim et al., 2019). We found that the transcriptional expression of Rbm47 were similar to that of Tjp1 α+ during preimplantation development. Immunofluorescence analysis revealed that Rbm47 protein was first detected in morula-stage embryos, and primarily localised to the nucleus in blastocyst embryos (Figure 1C and D).

To investigate the role of Tjp1 α+ in the stepwise formation of TJs during blastocyst development, we utilized electroporation to specifically knock down Rbm47 transcripts in the outer cells of 8-cell embryos. This approach was preferred due to its higher effectiveness and reduced degradation upto the blastocyst stage compared to the conventional method of siRNA microinjection into 1-cell zygotes. We knock down Rbm47 mRNA by up to ∼85%, which effectively suppressed Tjp1 α+ expression by up to ∼75% in blastocyst embryos. This is consistent with a previous study in human cell lines (Kim et al., 2019).

Interestingly, there were no significant differences in blastocyst development and TE permeability between the control group (84.62% and 6.42%, respectively) and the Rbm47 knockdown group (82.61% and 9.61%, respectively) (Figure 2 and Table 1). These findings suggest that the loss of Tjp1 α+ expression mediated by Rbm47 depletion did not impact the paracellular sealing of the TE and that Tjp1 α+ is not crucial for stepwise TJs assembly during blastocyst development.

**Table 1.** Preimplantation development and TJs integrity in Rbm47 knock down.

|  | Total<br>no. of<br>8-cell | Lysis | Intact<br>8-cell | Blastocyst | FITC<br>treated<br>Blastocyst | negative | positive |
| --- | --- | --- | --- | --- | --- | --- | --- |
| Control | 303 | 17 | 286 | 242<br>(84.62%) | 218 | 204 | 14 |
| KD | 297 | 21 | 276 | 228<br>(82.61%) | 229 | 207 | 22 |

Table 1.
|  | 8-cell | Lysis | Blasotycyst | (%) | FITC(-) | FITC(+) | Permeability |
| --- | --- | --- | --- | --- | --- | --- | --- |
| Control | 303 | 17 | 242 | 84.61 | 204 | 14 | 6.42% |
| KD | 297 | 21 | 228 | 82.62 | 207 | 22 | 9.61% |

**Figure 2.**
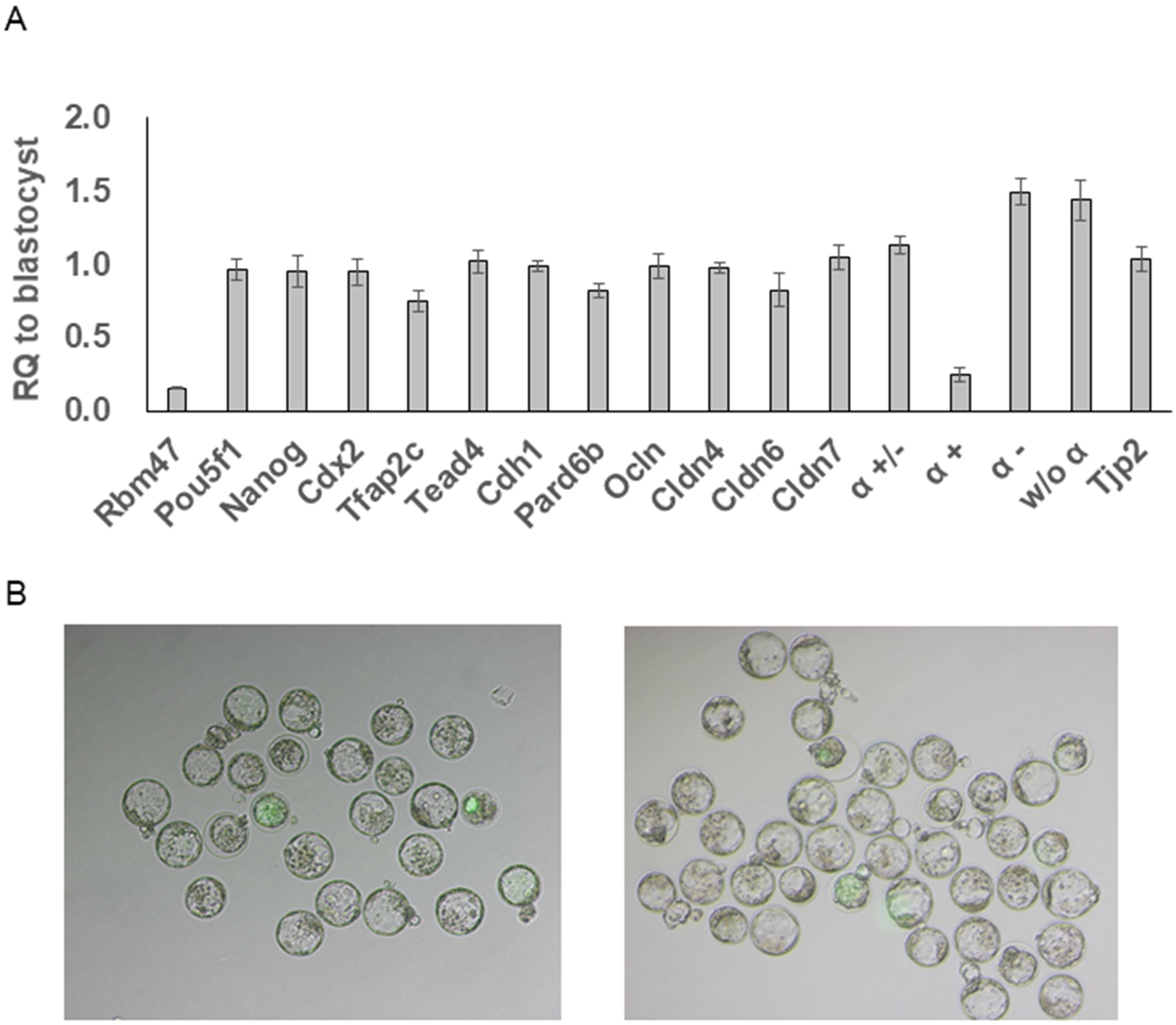
Effect of *Rbm47l* knockdown on gene expression and TJ permeability. (A) Transcript levels of genes associated with cell lineage, adherens junctions, tight junctions in Rbm47 KD blastocysts. (B) FITC-dextran uptake assay showed no significant differences in accumulation between control (left) and Rbm47 KD blastocysts (right), indicating no effect on tight junction permeability barrier. Relative quantification (RQ). Error bars represent mean ± standard error.

Moreover, qRT-PCR analysis in the KD blastocyst demonstrated that cell lineage (Pou5f1, Nanog, Cdx2, Tfap2c, Tead4), adheren junction (Cdh1), and tight junction-associated genes (Pard6b, Cldn4, Cldn6, Cldn7, Tjp1, and Tjp2) were not altered, except for Tjp1 α+ expression (Figure 1B). These gene expressions in the KD suggested that we may exclude multifunctional roles of Rbm47 such as RNA editing, and transcriptional activation in blastocyst development (Shivalingappa et al., 2021). We also confirmed that Rbm47 knockdown only affected the expression of Tjp1α+ inclusion by using another set of primers (w/o α; Figure 1A and 2A).

Our immunofluorescence analysis using variant-specific antibodies showed that loss of Rbm47 did not affect the expression of Tjp1 α-at the protein level, which is consistent with the transcript level. In control blastocysts, we observed two distinct continuous lines of Tjp1 α+/α-that overlapped at the apicolateral region of TE cells. However, in the Rbm47 knockdown blastocysts, we observed reduced expression of Tjp1 α+ with discontinuity and fragmented localisation of the apical region (Figure 3 A). These results suggest that that Tjp1 α+ may not play a significant role in culminating TJs assembly.

**Figure 3.**
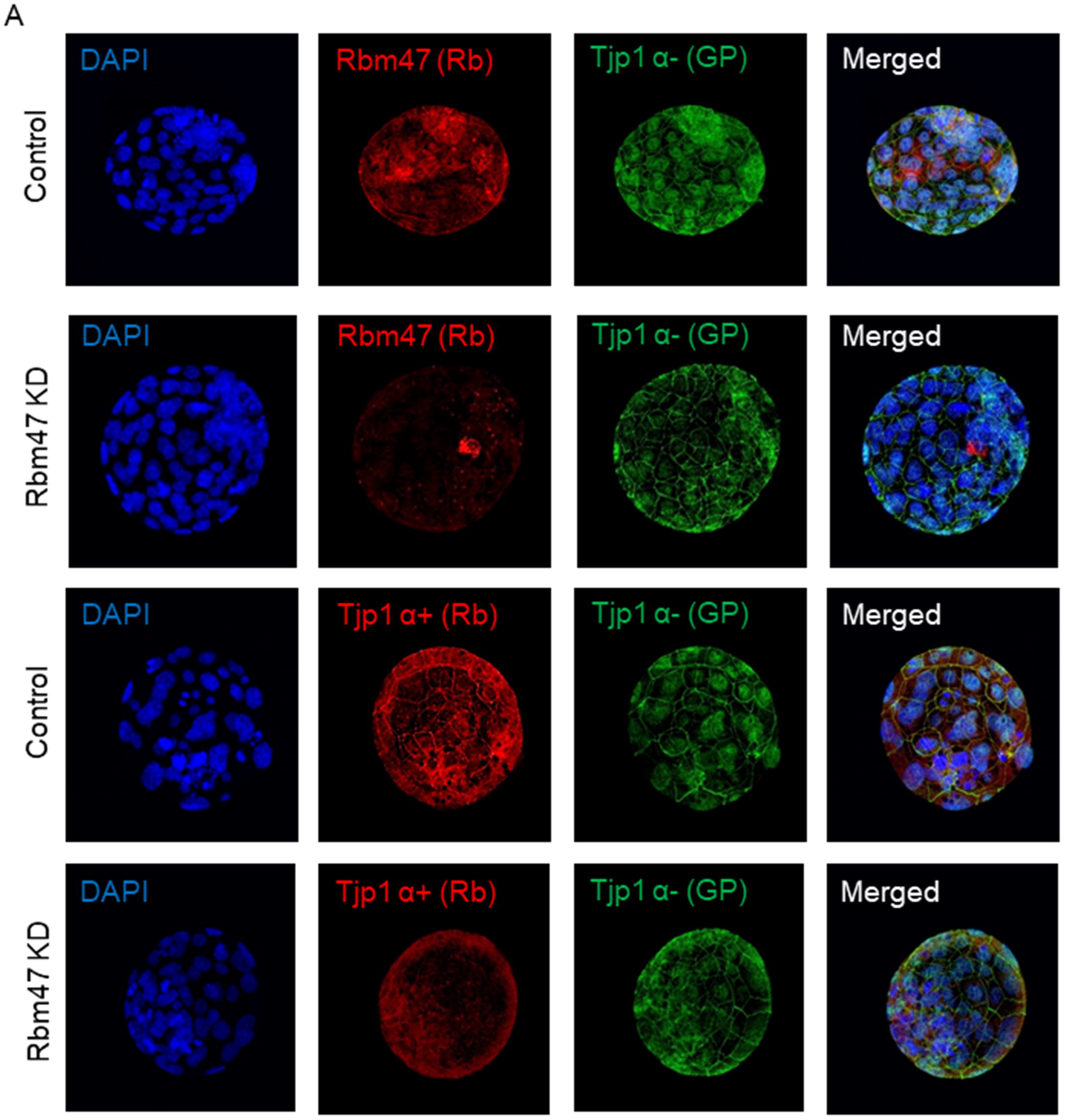
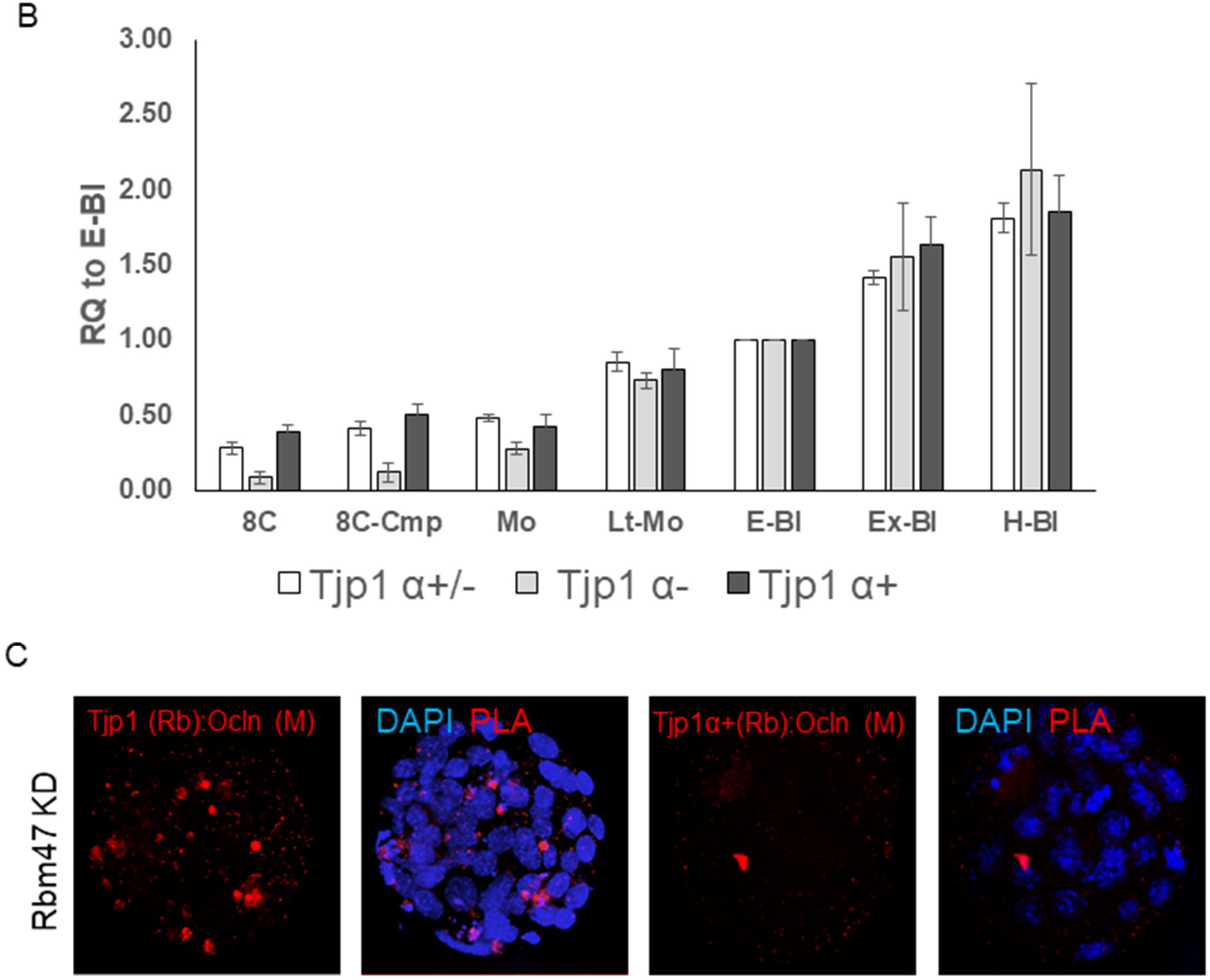
Effect of *Rbm47* knockdown on Tjp 1 expression and interaction with Ocln. (A) Expression and localisation of Tjp1 α+ and Tjp1 α-in the Rbm47 KD blastocyst. (B) Gene expression patterns of Tjp1 variants from 8-cell to hatching blastocyst (C) Proximity ligation assay (PLA) showing the interaction between Tjp1(mainly Tjp1 α-) and Ocln in blastocysts. 8C(8-cell), 8c-cmp (compacted 8-cell), Mo (Morula), Lt-Mo (Late morula), E-Bl (early blastocyst), Ex-Bl (Expanding-Bl), H-Bl (Hatching blastocyst). Antibodies host: Rabbit (Rb), Guinea pig (GP), mouse (m).

In this study, we indirectly reduced the expression of Tjp1 α+ variant by regulating alternative splicing, specifically depleting Rbm47 and targeting exon 20 (α-domain). As a result, the remainng expression level of Tjp1 α+ variant was relatively higher compared to direct depletion by siRNA, where the efficiencies were achieved around 90% (Choi et al., 2012; Jeong et al., 2022; Jeong et al., 2019). Additinally, expression of Tjp1 α+ variant isoform is lower than that of Tjp1 α-in preimplantation embryos (Sheth et al., 1997), suggesting that a minimal amount of Tjp1 α+ protein (∼25%) might be sufficient for the blastocoel formation. However, we postulated that continuous expression of Tjp1 α-during further blastocyst development may compensate for the role of Tjp1 α+.

Using qPCR, we measured the transcript levels of Tjp1 α+ and Tjp1 α-from 8-cell embryos to hatching blastocysts. Compared to the expression levels at the early blastocyst stage, Tjp1 α+ was increased by approximately 2.54 (RQ), and Tjp1 α-was increased by approximately 1.86 (RQ) (Figure 3B). This indicates that even after the formation of the blastocyst cavity, the expression of Tjp1 α-continues to increase, suggesting that Tjp1 α-is involved in the formation and maintenance of the cavity and can interact with other TJ proteins such as Ocln. To investigate whether Tjp1 α-could interact with Ocln in Rbm47 KD (Tjp1 α+ reduced) embryos, considering that the Tjp1 α+ variant has been reported to regulate the assembly of Ocln to the TJ membrane site (Sheth et al., 2000a; Sheth et al., 2000b), we conducted an in situ proximity ligation assay (PLA) using anti-Ocln and anti-Tjp1 instead of Tjp1 α-because probes for anti-Guinea pig IgG were unavailable for PLA. In Figure 3B, we observed an interaction between Tjp1 and Ocln proteins, localised at the apical regions of the blastomeres in the Rbm47 KD blastocyst. However, we did not observe any interaction between Tjp1 α+ and Ocln in the Rbm47 KD group. These findings suggest that main interactions with Ocln were from Tjp1 α-rather than Tjp1 α+ in the Rbm47 KD blastocyst due to the reduction of Tjp1 α+ expression.

In summary, we investigated the role of Tjp1 variant, Tjp1 α+, during mouse blastocyst development by using Rbm47 knockdown, a protein that regulates the alternative splicing of the α+ domain. We found that the reduction in Tjp1 α+ expression did not have a significant effect on blastocyst development or TE permeability. Our findings do not support the idea that Tjp1 α+ is necessary for the stepwise assembly of tight junctions during early development in mice.

## Materials and methods

All chemicals were purchased form Sigma-Aldrich (Sigma-Aldrich, St. Louis, MO, USA) unless otherwise stated.

### Embryo culture and knock down experiment

All animal manipulations were conducted in accordance with the guidelines set forth by the Institutional Animal Care and Use Committee of the Chungnam National University Animal Welfare and Ethical Review Body (License No. CNU-00702). Mouse embryos were collected and cultured as previously described (Jeong et al). In brief, fertilized 1-cell zygotes were collected from superovulated B6D2/F1 females mated with B6D2/F1 males (B6D2/F1; KOATECH, Pyeongtaek, Republic of Korea) and cultured in micro-drops of KSOM medium with 1/2 amino acids (EmbryoMax®; EMD Millipore, Billerica, MA) in a humidified atmosphere of 5% CO_2_ at 37°C until use.

To perform knockdown (KD) using electroporation, 8-cell stage embryos were washed three times with Opti-MEM I solution to remove serum. The embryos were then aligned within the electrode gap, which was filled with Opti-MEM I solution containing either 20 μM of *RBM47* siRNA or 20 μM of scrambled siRNA (SMART pool siGenome; Dharmacon, Lafayette, CO, USA). Electroporation was carried out using Genome Editor TM (BEX, Tokyo, Japan) with 25 V (3 msec ON + 97 msec OFF) for 7 cycles. Following electroporation, the embryos were immediately retrieved from the electrode chamber and rinsed four times in M2 medium, followed by two rinses with culture medium. Subsequently, the embryos were returned to the culture medium (Hashimoto and Takemoto, 2015; Kaneko et al., 2014)..

### Quantitative real-time PCR (qRT-PCR)

To isolate total RNA, embryos at each stage (one-, two-, four-, and eight-cell, morula and blastocyst) were collected and processed using a PicoPure RNA Isolation Kit (Arcturus, Mountain View, CA, USA). The isolated RNA was then reverse transcribed into complementary DNA (cDNA) using oligo(dT) and reverse transcriptase (SuperScript II, Invitrogen, Carlsbad, CA, USA). Real-time PCR was performed on a StepOne Plus Real-Time PCR System (Applied Biosystems, Foster, CA, USA) using SFCgreen® (BIOFACT, Daejeon, Republic of Korea) as the fluorescent probe. Gene-specific primers were used, including Rbm47 (forward: 5’-CAGCTTGTTTCCTGCTACGC -3’, reverse: 5’-CTCTGAGGCAAACCCGAACT-3’), Tjp1 (α+/α-) (forward: 5’-AGATAGCCCTGCAGCCAAAG -3’, reverse: 5’-AGGACGGCCTCTTCCCTTAT -3’), Tjp1 (α+) (forward: 5’-GCTCACGTAGTGCTCAGAGG -3’, reverse: 5’-CAGAATACGGCTCCTTCCTGT -3’), Tjp1 (α-) (forward: 5’-CCTGACGGTTGGTCTTTTGC -3’, reverse: 5’-ACAGTTGGCTCCAACAAGGT-3’), and Tjp1 (without α) (forward: 5’-CAGCCTCTCAACAGGTGTA-3’, reverse: 5’-TGGGTGACCAAGAGCTG-3’). Additional primer sequences can be found in (Choi et al., 2012). For relative quantification of gene expression between control and KD embryos, we used the 2^−ΔΔ^Ct method was with *Ubtf1* as an endogenous reference gene. To normalize samples from different stages of preimplantation embryos, *Gfp* was spiked into the same number of each embryo prior to RNA isolation and used as an exogenous reference.

### Immunocytochemistry (ICC) and Proximity ligation assay (PLA)

Immunocytochemistry (ICC) and proximity ligation assay (PLA) analysis of preimplantation embryos were conducted following previously described methods (Jeong et al., 2022)(Jeong et al., 2022). Briefly, embryos were fixed with 3.7% paraformaldehyde in phosphateLJbuffered saline (PBS) for 20 min, permeabilized with 0.1% (v/v) Tween-20 in PBS, and blocked with 0.1% (v/v) bovine serum albumin (BSA) in PBS for 1 h at room temperature. Primary antibodies against RBM47 (HPA006347, Sigma-Aldrich), TJP1 α+ (HP9044; Hycult Biotechnology, Uden, Netherlands) and TJP1 α-(HP9045; Hycult Biotechnology) were diluted and incubated with the embryos overnight at 4°C. The embryos were then incubated with Alexa Fluor 488 and 546 secondary antibodies (Molecular Probes, Eugene, OR, USA) for 1 h at room temperature.

For the PLA assay, embryos treated with a pair of primary antibodies such as TJP1 (40-2200, Invitrogen, Carlsbad, CA, USA), and Ocln (OC-3F10, Invitrogen), and TJP1 α+ and TJP1 were incubated with PLA probes (anti-mouse IgG and anti-rabbit IgG), followed by ligation and amplification according to the manufacturer’s protocol (Duolink® In situ Red Starter kit, Sigma-Aldrich). The embryos were then mounted in ProLongTM Diamond Antifade Mountant with DAPI (Invitrogen), washed three times, and visualized using a fluorescence laser confocal microscope (K1-FluoRT, Nanoscope system, Daejeon, Republic of Korea). All images were analyzed using ImageJ 1.53e (http://imagej.nih.gov/ij).

### TJ permeability

Control and Rbm47 KD embryos were cultured until 120 hours post-hatching (hph). Subsequently, blastocysts were incubated for 10 minutes in modified KSOM medium supplemented with 1 mg/mL 4 kDa FITC-dextran. After incubation, the blastocysts were washed, and the accumulation and position of FITC-dextran in the blastocoel was confirmed under an inverted fluorescence microscope with the bright-field mode (Eclipse Ti-U, Nikon).

### Statistical analysis

Data were analysed by Student’s *t*-test using GraphPad Prism (Version 5.03, GraphPad Software, San Diego, CA, USA), and are presented as mean ± standard error. *P* values < 0.05 were considered significant unless otherwise stated

## Author contribution

IC conceived and designed the study. I.C. and J. J. performed animal and experiments. IC prepared the manuscript. All authors read and approved the final manuscript.

